# Low-cost, sub-micron resolution, wide-field computational microscopy using opensource hardware

**DOI:** 10.1101/460055

**Authors:** Tomas Aidukas, Regina Eckert, Andrew R. Harvey, Laura Waller, Pavan C. Konda

## Abstract

The revolution in low-cost consumer photography and computation provides fertile opportunity for a disruptive reduction in the cost of biomedical imaging. Conventional approaches to low-cost microscopy are fundamentally restricted, however, to modest field of view (FOV) and/or resolution. We report a low-cost microscopy technique, implemented with a *Raspberry Pi* single-board computer and color camera combined with Fourier ptychography (FP), to computationally construct 25-megapixel images with sub-micron resolution. New image-construction techniques were developed to enable the use of the low-cost Bayer color sensor, to compensate for the highly aberrated re-used camera lens and to compensate for misalignments associated with the 3D-printed microscope structure. This high ratio of performance to cost is of particular interest to high-throughput microscopy applications, ranging from drug discovery and digital pathology to health screening in low-income countries. 3D models and assembly instructions of our microscope are made available for open source use.

## Introduction

Low-cost, high-performance portable microscopes are essential tools for disease diagnosis in remote and resource-limited communities [1]. A fundamental requirement is to combine wide field of view (FOV) with the high resolution necessary for imaging of sub-cellular features of biological samples. This underpins efficient inspection of extended, statistically-significant areas for screening of, for example, cancer, malaria, or sickle cell anemia [2]. In conventional imaging, the number of pixels in the detector array constitutes a hard limit on the space-bandwidth product (SBP – the number of pixels in a Nyquist-sampled image) [3,4] so that increased FOV can be achieved only at the expense of reduced spatial resolution. SBP can be increased using larger detector arrays coupled with higher-performance, wide-field aberration-corrected optics, or by mechanical scanning, but these approaches add complexity, cost and bulk [5,6].

Several low-cost portable microscopes have been proposed [7–12], but they all suffer from the problem of small SBP. Early progress towards low-cost microscopy has involved the use of a high-cost microscope objective lens coupled to a mobile-phone camera [7] and such instruments tend to suffer from a higher system cost, vignetting, short working distance, small depth of field (DOF) and narrow FOV. Lower-cost implementations have been reported in which the microscope objective is replaced by a camera lens from a mobile phone [8], or a ball lens [9], but their resolving power is limited by the small numerical aperture (NA) and high aberrations. Of these implementations, the use of mobile-phone camera lenses as objectives places an upper limit on the SBP: for example a 4-μm spatial resolution across 9mm^2^ FOV corresponding a SBP of 2.25Mpixel [8]. The 4-μm resolution is insufficient for observing sub-cellular features and while a higher NA can be obtained using ball lenses, providing a resolution around 1.5 μm, they suffer from small SBP [8,13].

We report a low-cost, wide-field, high-resolution Fourier-ptychographic microscope (FPM) [14], implemented with 3D-printed opto-mechanics and a *Raspberry Pi* single-board computer for data acquisition as shown in Fig. 1(a). High-SBP images are constructed from multiple low-resolution, detector-SBP limited images, captured in time-sequence using oblique illumination angles yielding a SBP that is much greater than that of the detector. We demonstrate 25-Megapixel microscopy using a 4-Megapixel detector array. The tilted illuminations provide translations of higher spatial-frequency bands into the passband of the objective lens [15]. Stitching of images in the frequency domain is implemented using an iterative phase-retrieval algorithm to recover high-resolution amplitude and phase of the sample image [16,17], as well as aberrations due to the objective [14]. Recovery of phase information enables imaging of unstained transparent samples [18] and computational calibration of illumination angles during image reconstruction is able to correct errors arising from misalignment of various components [19,20], which is of particular importance for microscopy using low-cost 3D-printed devices.

**Figure 1.**
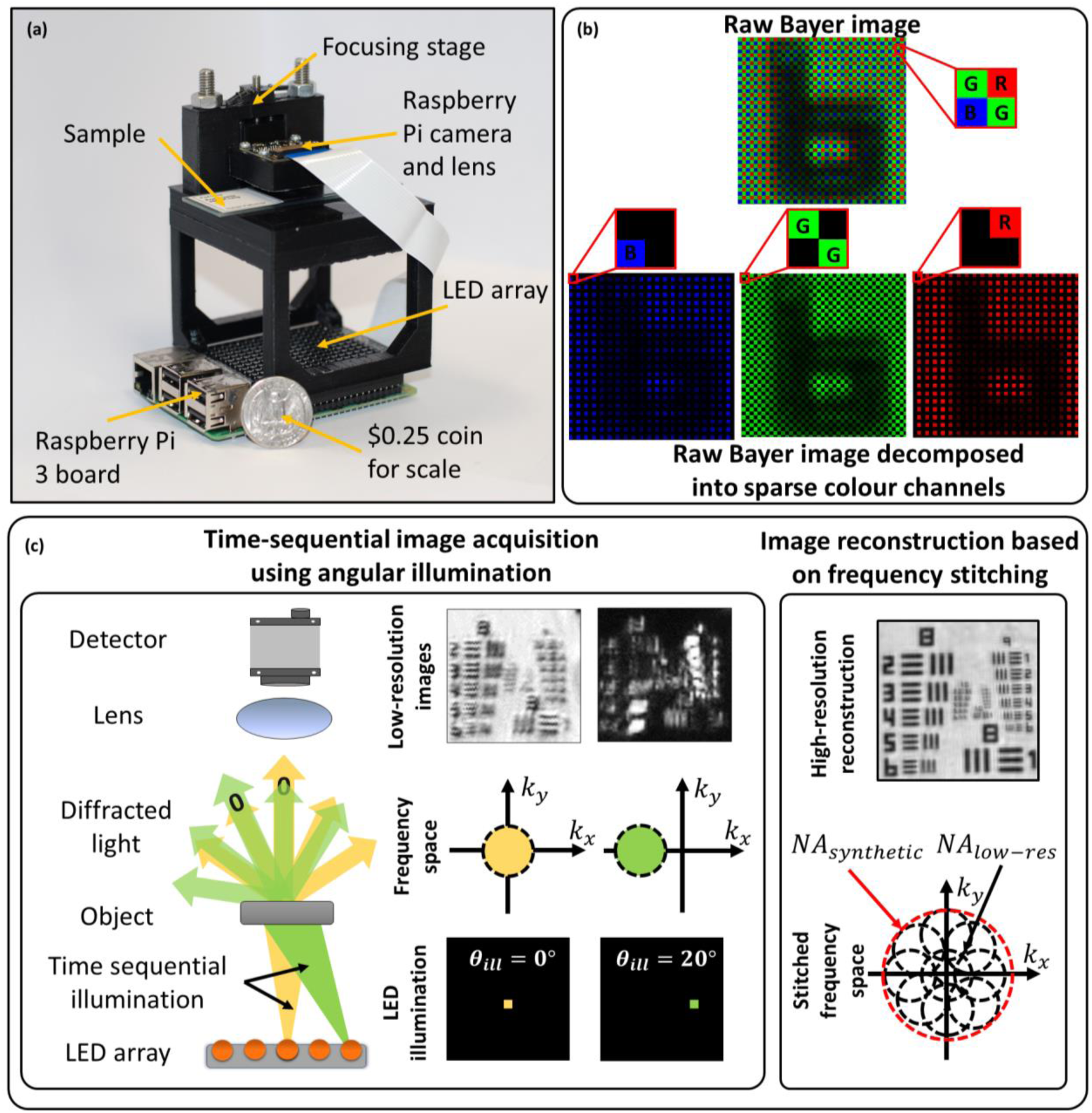
**(a)** Experimental setup next to a quarter US dollar for scale. *Raspberry Pi 3* single-board computer board (placed at the bottom) enables wireless image acquisition and data transfer without the need for a PC. **(b)** Bayer color filter array indicating RGGB pixel arrangement. (c) In FPM several low-resolution images are obtained in time sequence, each illuminated with a corresponding to the object illuminated from a different angle. Angular diversity enables to obtain multiple frequency regions, which can be stitched together into a single high-resolution, wide-field image.

In previous demonstrations of a low-cost 3D-printed FPM, the SBP was limited by the severe off-axis aberrations of the mobile-phone camera lens (1.5 μm resolution across 0.88mm^2^ FOV giving a SBP of 1.56Mpixels), and employed a science-grade, high-cost monochrome sensor [21]. Exploiting the mass market for consumer color sensors in mobile phone cameras, we demonstrate the first use of a low-cost consumer color camera in FPM, to gain more than an order-of-magnitude cost reduction for an equivalent SBP. The main difference between the two sensor types is the spatial-spectral filtering provided by the Bayer filter array, which encodes recorded images into sparse red, green, and blue channels. While the decoding processes follows a standard demosaicing procedure (individual RGB channels are interpolated and stacked into a 3D matrix), the loss in image information due to sparse sampling requires special treatment within the FPM reconstruction algorithm. We address the sparse sampling problem and present new robust algorithms for calibrating the 3D printed system for high-quality image reconstruction. In addition, the *Raspberry Pi* single-board computer used for controlling the camera and illumination LEDs performs autonomous data acquisition, providing portability and compactness, such as is required for use inside incubation systems.

In the next section, simulations to study the impact of the Bayer filter array and the experimental results from our system are presented. Implications of the results and future directions are discussed in the later sections. The methods section includes descriptions of the experimental setup, data-acquisition, data processing and calibration procedures. We also include the necessary CAD files and an instruction set to build the FPM presented in this article (supplementary material S1).

## Results

The *Raspberry Pi* camera (a low-cost device that complements the *Raspberry Pi* computer) employs a low-cost CMOS sensor, such as is typically found in mobile phones. It employs a Bayer filter (red, green and blue filters arranged on a 2D matrix in a 2×2 RGGB pattern [22] (Fig. 1(b))). This divides pixels on the sensor between the three color-filters resulting in sparsely sampled images: red channel – 75% empty pixels, green channel – 50% empty pixels and blue channel – 75% empty pixels. The empty pixels are demosaiced (using bilinear interpolation) to produce a perceptually acceptable photographic image.

In FPM, the reconstruction algorithm [18] (see Methods) involves a step to iteratively recover amplitude and phase of the low-resolution images, where the estimated amplitude is replaced by the experimentally obtained images. In color cameras, the experimental image has empty pixels (due to the Bayer filter) whose values are unknown. We have considered two approaches for mitigation of the sparse sampling due to the Bayer filter. The first, a *sparsely-sampled reconstruction* (SSR) algorithm [23], updates only the non-empty image pixels, relying on the FPM reconstruction to estimate the empty image pixels. This approach increases the number of unknowns in the system and can have slower convergence or failure to converge. In a second approach, the empty pixels are estimated instead from demosaicing enabling the use of a conventional FPM recovery; we refer to this approach as *demosaiced reconstruction* (DR). With DR the interpolation errors introduced in demosaicing can introduce artefacts in the reconstruction. We report below a comparison of image-recovery accuracy using SSR and DR recovery applied to simulated data.

Convergence of the FPM reconstruction algorithms requires the experimental design conditions to satisfy Nyquist sampling of the image by the detector array and to have approximately 50% overlap between the frequency bands selected by adjacent illumination angles (Fig. 2(c2)) [24]. We assess here using simulations, how these requirements are modified by the reduced sampling rate associated with the sparse sampling of the Bayer matrix. Image quality is compared to recovery from non-Bayer-encoded images.

**Figure 2.**
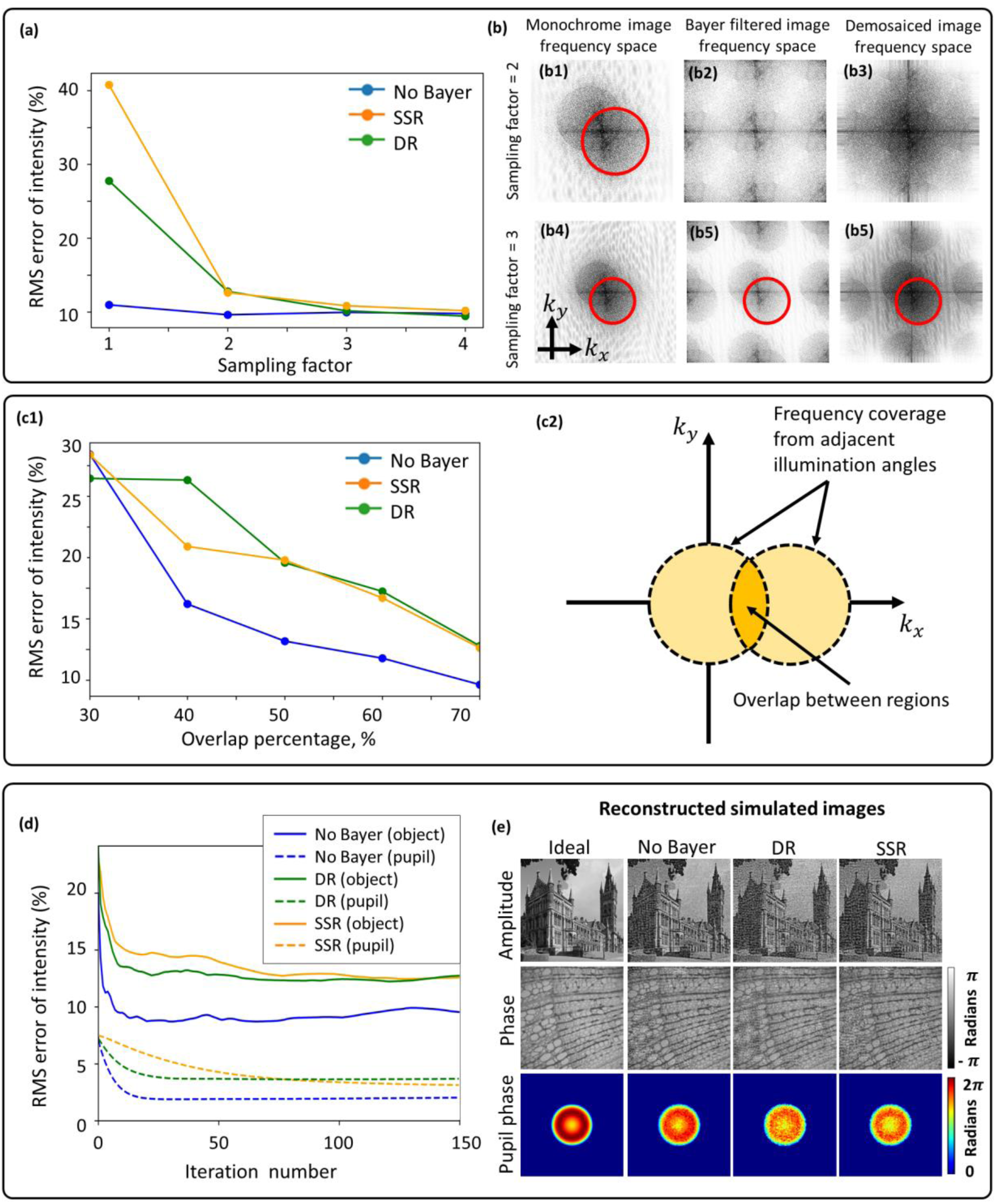
**(a)** Demosaiced and sparsely-sampled reconstruction accuracy for different sampling factors showing that a factor of two is required when using DR and SSR methods; 70% overlap area in the frequency domain. **(b)** Frequency spectra of monochrome and color sensor images showing frequency replicates introduced by the Bayer filter and how it distorts the circular boundary. The boundary becomes undistorted only for a sampling factor of 3. **(c)** Demosaiced and sparse reconstruction accuracy for different frequency overlap percentages together with a diagram explaining what is meant by the overlap percentage between adjacent frequency regions. As expected, accuracy improves as overlap increases. **(d)** Reconstruction convergence plots for object amplitude and pupil phase (70% overlap and sampling factor of 2), indicating better performance of demosaiced reconstruction. **(e)** Reconstructed simulated images.

Using the far-field approximation [15], the image intensity for a color channel can be written as

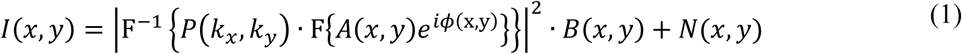

where (*k_x_*, *k_y_*) are coordinates in frequency space, (*x,y*) are coordinates in real space, *P* is the pupil function, *A* and *ϕ* are the amplitude and phase distributions of the input object respectively, *B* is a binary mask corresponding to the color channel’s filter arrangement on the RGGB Bayer matrix, *N* is the added Gaussian image noise and F is the Fourier transform operator. Since robustness of the reconstruction is strongly dependent on the aberrations present in the pupil plane, they were simulated by including defocus and spherical optical aberrations generated using Zernike polynomials. We employed the Root-Mean-Squared (RMS) error between high-resolution reconstructed image and the expected ideal simulated image as a metric of image quality. We employed 150 iterations, which was more than sufficient for the FPM algorithms to converge.

In an imaging system, the image-sampling frequency is defined as *f_sampling_* = *M/PS*, where *M* is the magnification and *PS* is the pixel size. This sampling frequency must satisfy the Nyquist sampling criterion, defined as twice the optical cut-off frequency, to avoid aliasing:

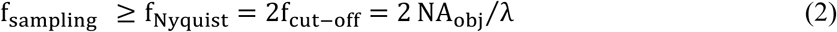

The image-sampling frequency can be controlled in the experimental design by modifying the magnification since the pixel size is fixed by the camera sensor characteristics. To achieve the widest FOV possible without aliasing, the sampling factor (*f_sampling_*/*f_Nyquist_*) must be unity. For Bayer sensors, intuitively the effective pixel width is 2× larger due to the empty pixels in each color channel of the Bayer filter array, hence, the magnification needs to be increased by a factor of two compared to a monochrome detector array to compensate, i.e., the required sampling factor will be two. Since increasing the magnification reduces the FOV, simulations were performed (Fig. 2 (b)) to assess whether the FPM reconstruction methods could converge with under-sampling to achieve the highest SBP.

### Comparison of FPM reconstruction techniques for Bayer images

Sparsely-sampled reconstruction has been shown to be effective for aliased images with 75% sparsity [23], offering an advantage in terms of maximum achievable SBP compared to demosaiced reconstruction. However, as can be seen in Fig. 2(a), the image reconstruction from Nyquist-sampled Bayer images exhibited large RMS errors of 20-30% compared to 10% for non-Bayer images. Reduced image quality for reconstruction from Bayer-sampled images is expected due to aliasing artefacts; however, these findings differ from the conclusions in [23]: probably due to practical differences in implementation, which did not involve compensation of optical aberrations and benefitted from low-noise data recorded by science-grade cameras. This enabled reconstruction of high-resolution images from data with 75% sparsity. However, in our implementation recovering the system aberrations and dealing with the detector read noise is crucial, hence, both SSR and DR reconstruction methods require an additional 2× magnification to satisfy the Nyquist sampling criterion.

In Fig. 2(c1), the requirement for overlap between the spatial frequencies recorded by two adjacent LEDs is assessed. It suggests that RMS errors for SSR start to converge at ∼40% overlap compared to 50% for DR; this is in agreement with the requirements for non-Bayer sensors [24]. Since the additional 2× magnification is used in these simulations, the frequency overlap requirement achieved is similar to the requirement for non-Bayer systems. Using these two optimal system parameters (2× additional magnification and a 70% frequency overlap), the overall convergence for DR and SSR and non-Bayer systems is compared in Fig. 2 (d). It can be observed that DR has better convergence and pupil recovery than SSR. The RMS errors in the final reconstructions are close for DR and SSR, hence it can be concluded that DR has better convergence properties despite both reconstruction techniques resulting in similar reconstruction quality. All reconstructed images are shown in the supplementary material S2.

### Algorithmic self-calibration of LED misalignment

Our system is implemented using 3D printed components and intended to be portable; hence, it may become easily misaligned, affecting primarily the illumination angles (LED positions). In addition, image distortion and field curvature change the relative LED positions distinctly across the FOV [25]. We have implemented a recently-developed self-calibration algorithm for LED position misalignment correction [20], solving the issues of image distortion and misaligned components with relatively good computational efficiency (see Methods). In this algorithm the intensity image of an off-axis illuminated brightfield image is Fourier transformed to produce two overlapping circles, centered around the illumination direction. Using image processing techniques, we can find centers of these circles providing a better calibration for the LED positions; hence, the calibration accuracy depends on how well these circles are delineated.

While a sampling factor of two is sufficient (for a monochrome sensor), our simulations suggest (Fig. 2(b)) that artefacts introduced by the Bayer matrix require the sampling factor to be around three to produce an undistorted circular boundary, regardless of demosaicing. The Bayer pattern can be treated as a periodic grating; hence, it produces frequency replicas (similar to diffraction orders), a type of aliasing artefact, which distort circle boundaries indicated by Fig. 2(b). Hence, by increasing the sampling frequency, the separation between these frequency replicas is increased to preserve the boundaries. In practice, the change in illumination wavelength varies the sampling factor for a fixed magnification since the sampling frequency is fixed but the Nyquist frequency changes; hence, 3× sampling factor requirement for red (630nm) (enough for calibrating LED positions) results in 2× sampling factor in blue, the minimum required for overcoming Bayer sampling. This suggests that the red channel can be used for LED position calibration without losing additional SBP due to the increased sampling requirement. The FOV is divided into several segments and processed independently in FPM, hence the distortion is tackled by calculating the relative LED positions for each of these segments independently (see methods for the recovered distortion of the system).

### Experimental results

Our FPM device (Fig. 1(a)) achieves high performance at low cost by use of mass-produced consumer electronics: a conventional mobile-phone-type color camera (with the lens displaced from the normal infinite-conjugate imaging position to enable short-range imaging), a *Raspberry Pi* single-board computer for data acquisition and an off-the-shelf LED array (*Pimoroni Unicorn Hat HD*) for synthesis of a programmable illumination that enables synthesis of a higher NA. The total component cost is about $150, but mass production of such a device would further reduce the component cost. The lens from the *Raspberry Pi Camera v2.0* provides 0.15NA and 1.5× magnification when placed 7mm from the object. A 16×16 array of LEDs with 3.3-mm pitch was located 60 mm below the object providing 0.4-NA illumination to enable synthesis of 0.55-NA FPM images. The FPM yields a 25-megapixel SBP: that is 870-nm resolution (*NA* = 0.55) – sufficient for sub-cellular imaging across a 4mm^2^ FOV. FPM also enables multiple imaging modalities, including phase-contrast and darkfield imaging, combined with extended DOF and computational aberration correction [26,27].

Computational correction of errors due to imperfect calibration (such as component misalignment and aberrations) is highly dependent on image quality, which is compromised by the Bayer matrix due to optical attenuation and spectral overlap and spectral leakage between the RGB channels. While signal-to-noise ratio was maximized by independent optimization of integration times for each illumination angle, the spectral overlap of the Bayer spectral filters was mitigated by each red, green and blue LED in a time sequence rather than simultaneously.

We used a standard USAF resolution test chart (Fig. 3(a)) to quantitively assess the performance and resolution improvement. Analysis of the reconstructed images shows a resolution improvement from group 8 element 4 (Fig. 3(a3)) to group 10 element 3 (Fig. 3(b6)) (using 470nm (blue LED) illumination), which corresponds to a three-fold resolution improvement from 2.8μm (incoherent-sum) to 780nm. This resolution improvement is the result of the large synthetic NA offered by FPM, which is defined as *NA_FPM_* = *NA_ill_*+*NA_objective_*. Experimental results agree with the theoretical predictions, which give an increase in NA from *NA_coherent_* = 0.15 (*NA_ill_* = 0, *NA_objective_* = 0.15) to *NA_FPM_* = 0.55 (*NA_ill_* = 0. 4, *NA_objective_* = 0.15). While reconstruction quality shown in Fig. 3(c1-c9, b1-b9) is nearly identical for both the DR and SSR, the DR offers faster convergence, since the SSR needs to iteratively recover the missing pixels that are readily available through demosaicing in DR. The impact of spectral overlap was demonstrated by illuminating the sample using RGB LEDs simultaneously (white light) and reconstructing each color channel. Artefactual reconstructions (Fig. 3(d1-d3)) are a result of the broken assumption of monochromatic light that is implicit in FPM and could be mitigated by a spectral multiplexing algorithm [28].

**Figure 3.**
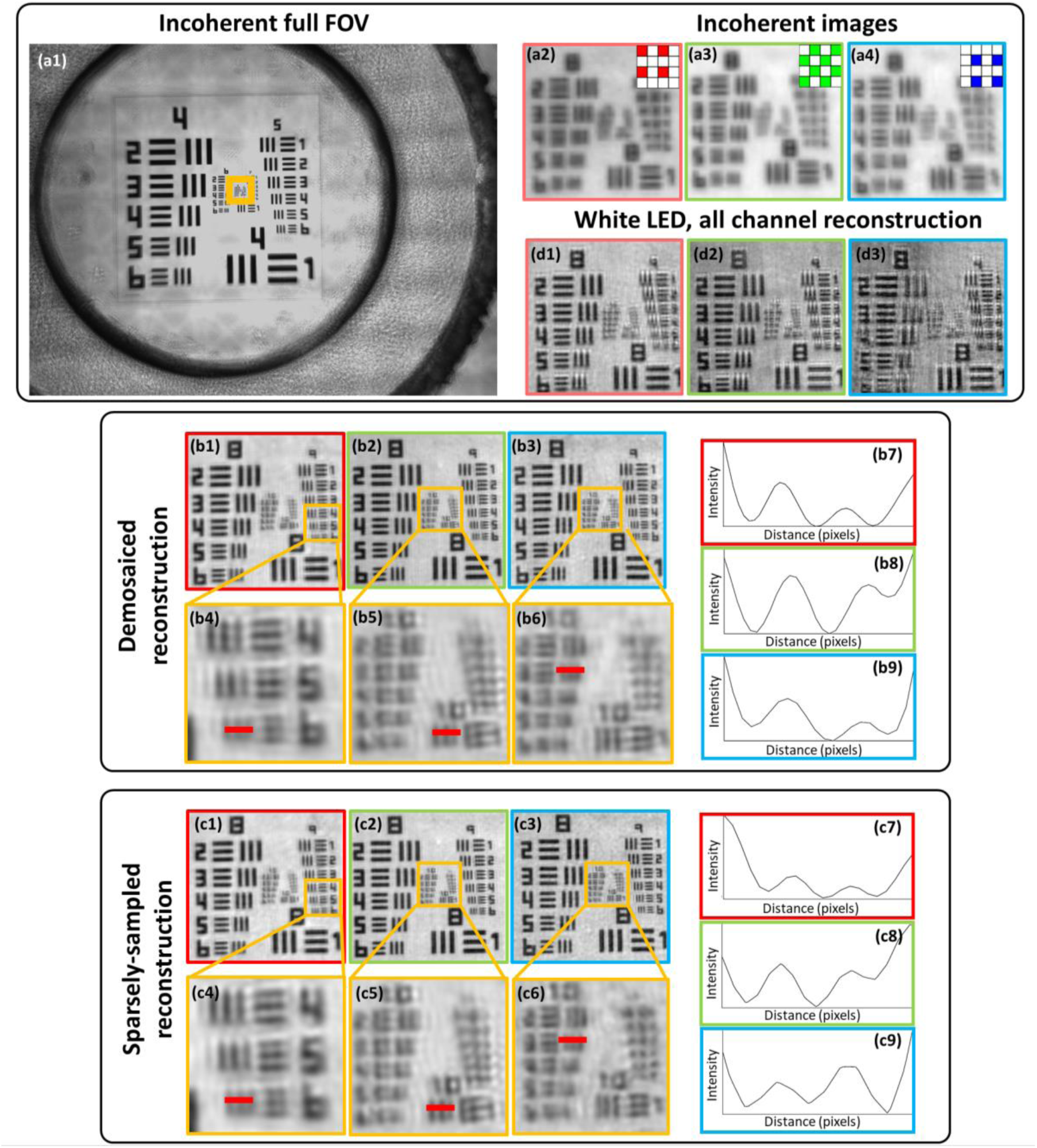
Reconstructions of a USAF resolution chart. (**a1-a4**) Incoherent raw images. (**b1-b9**) Demosaiced reconstructions and (**c1-c9**) sparsely-sampled reconstructions together with line profiles of the smallest resolved USAF target bars. The maximum achieved resolution using the blue LED was 780nm based on group 10 element 3. (d1-d3) Reconstructed images with RGB LEDs used in parallel for illumination demonstrating the reduced reconstruction quality due to the spectral overlap between the color channels. The respective color channels are indicated by the red, green and blue borders of the left, middle and right images

Lastly, we have demonstrated experimentally that our reconstruction algorithms can compensate for high-levels of optical aberrations associated with the simple low-cost objective lens. Reconstructed images of a lung carcinoma (Fig. 4(a,b)) show high-quality reconstruction across the full FOV despite the presence of off-axis aberrations, which are recovered and corrected within the reconstruction procedure without requiring additional data. It can be observed clearly in Fig. 4(d1), that the raw image is severely aberrated compared to (c1), but the reconstruction (d2) is of similar quality to the central FOV section (c2). The phase images shown in Fig. 4(c3, d3) demonstrate the capability of imaging unstained samples. It can be seen from Fig. 4(a) that without aberration correction the FOV is limited by aberrations to a central area of ∼1mm^2^ while the FPM correction of imaging aberrations increases the usable area of the FOV by a factor of four.

**Figure 4.**
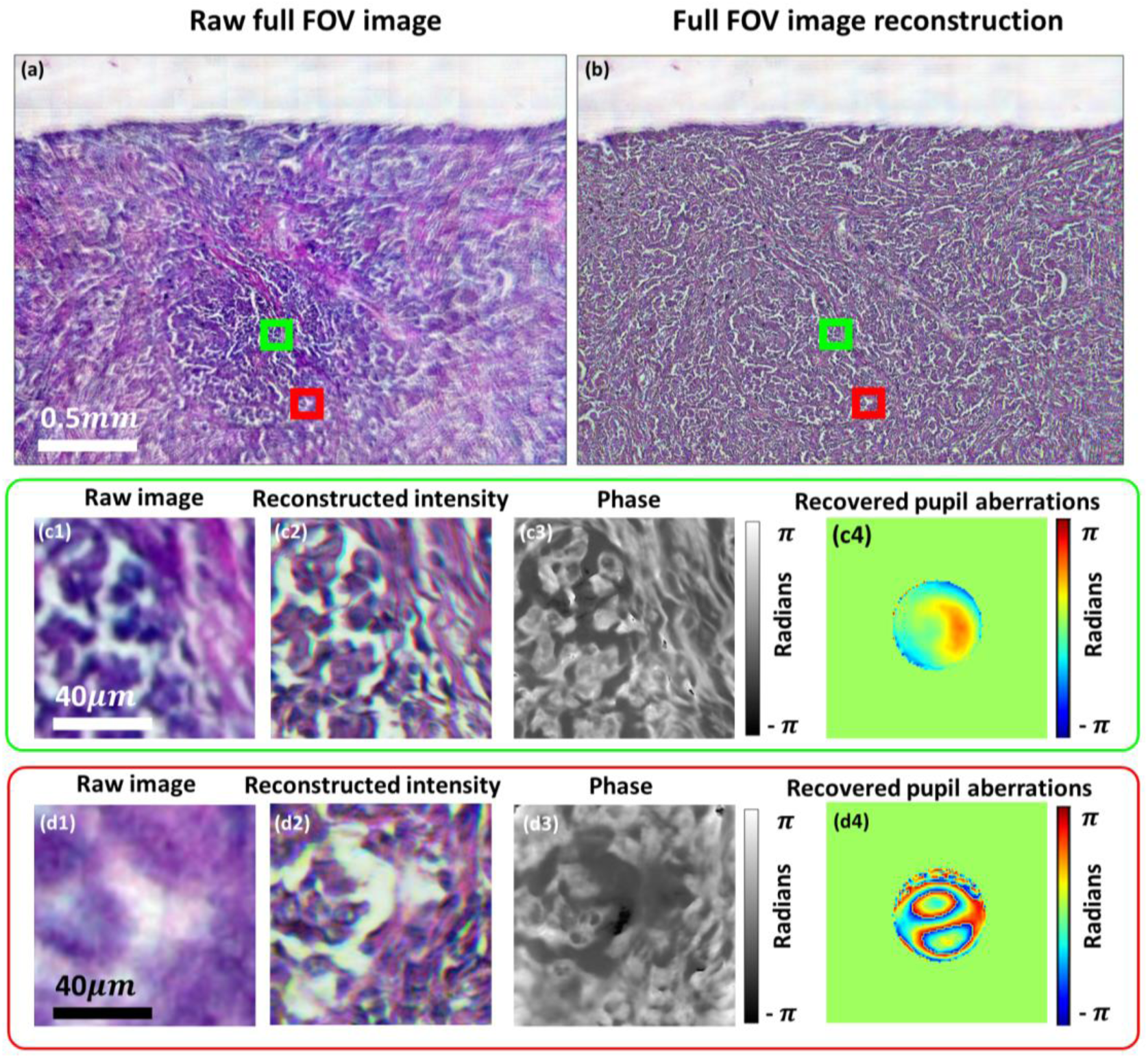
(**a**) Reconstructed and (**b**) raw lung carcinoma images. (**c1, d1**) are the captured raw, low-resolution images and (**c2-c3, d2-d3**) intensity and phase reconstructions for two different segments of the FOV. (**c4, d4**) Recovered pupils with aberrations.

## Discussion and conclusion

We have described the first demonstration of low-cost FPM, enabled by implementation using consumer-grade color cameras. We achieved a 4mm^2^ FOV and 780nm resolution (*NA* = 0.55), giving 25-megapixels SBP recovered from 256, 8-megapixel images. Compared to previous reports of low-cost, mobile microscopes [8] the resolution of our system is a factor of 5 times better with the added advantage of a 4-times longer working distance (due to the low-NA lenses). Compared to systems where mobile-phone cameras are equipped with expensive microscope objectives [12] our microscope offers 100-fold larger FOV without sacrificing resolution. Compared to a previously demonstrated 3D-printed FPM [21], we report an increase in the FOV area by a factor 5 and resolution by almost a factor of 2, while the use of a color sensor instead of a more specialist monochrome sensor reduced the cost by 1-2 orders of magnitude. The improved performance of our system is made possible by improved aberration correction and calibration strategies capable of coping with simple, low-cost components [23]. It should be noted that (1) due to the additional magnification required by the Bayer filtering, the effective SBP achieved from each 8-megapixel image is only 2 megapixels and (2) the 25 megapixels SBP corresponds to the number of pixels in the image, but each pixel in the reconstructed image contains both amplitude and phase information. Although the recording of 256 images may seem a high number, this degree of redundancy is typical and necessary with FPM [14], but can be reduced by a factor of up to 10 by using illumination multiplexing [29].

Our stand-alone microscope weighs only 200 grams and has external dimensions of 6cm x 9cm x 11cm. Data acquisition is autonomous offering major cost-savings and is ideal for applications such as cell-culture studies or point-of-care-testing applications that require field-portable devices. The *Raspberry Pi 3* computer-board enables wireless image acquisition, data transfer, and has potential for on-board FPM-based image reconstruction. Since image reconstruction is currently a computationally-intensive process we transferred the data to an external PC for processing, but in practice it would be possible to transfer the data onto a server network to perform the computations. Also, the use of a trained neural network for image recovery has been shown to improve image reconstruction speed by up to 50 times [30], which is particularly attractive for systems with lower computational power. However, neural network use for medical applications requires an investigation into the availability of training datasets, or data overfitting [31].

One major shortcoming of FPM is the time-sequential data acquisition, but image acquisition time of <1s has been demonstrated in FPM using LED multiplexing [18,29], which offers potential for an order-of-magnitude improvement in imaging speed. Techniques such as multi-aperture Fourier ptychography [32,33] can further increase throughput, as is essential for fast biological processes. Reduced image-acquisition times are also facilitated by replacement of the planar LED array with a dome-shaped array [34], where all LEDs are oriented towards the sample offering improved illumination efficiency. Lastly a further factor of 6 increase in data acquisition speeds could be achieved by removing the high latency introduced during sequential read-out of our CMOS cameras.

We have demonstrated that Fourier ptychography can be performed by using low-cost commercial-grade Bayer color sensors, off-the-shelf consumer components and 3D-printed parts. This is enabled by robust pre-processing and reconstruction strategies. Moreover, we used a *Raspberry Pi 3* single-board computer for image acquisition and image transfer. The result is a highly compact, stand-alone microscope, with a component cost of $150, that is capable of wide-FOV, high-resolution imaging. The proposed microscope is suitable for cell-culture studies (its compactness enables it to fit inside an incubating chamber) and point-of-care diagnostics. Due to the simplicity of our setup, it is suitable for use as a teaching tool for computational optics in undergraduate labs and in research labs for conducting further research in FPM.

## Methods

### Experimental setup

Instructions for construction of our microscope shown in Fig. 1(a) can be found in supplementary material S1. To minimize the cost of our microscope we used easily accessible off-the-shelf, low-cost components. We chose a finite-conjugate microscope design because it requires only a single lens. Sample and focusing stages were custom designed and 3D-printed using a *Ultimaker 2+* 3D printer. A *Raspberry Pi V2 NOIR* camera module was used (8 megapixels, 1.12um pixel size) which contains a 3-mm focal-length camera lens, which was remounted and displaced from the sensor to achieve ∼1.5× magnification. Frequency overlap of ∼70% was obtained by placing the *Unicorn HAT HD* 16×16 LED array (3.3mm pitch) 60mm below the sample stage. The RGB LED array has peak illumination wavelengths of 623nm, 530nm, and 470nm. The low-resolution microscope has 0.15 NA (providing 5-μm resolution at 470nm), 2.42×1.64mm^2^ FOV, and a 7-mm working distance. The synthetic NA achieved after FPM reconstruction was 0.55. Since the lens is used away from the intended infinite-conjugate position, the aberrations become progressively more severe toward the edges of the FOV. This could be mitigated be use of two back-to-back, co-aligned lenses [8] with the penalty of reduced working distance and added experimental complexity.

### Data acquisition

Experimental low-resolution images were obtained using all 256 LEDs in the LED array. The *Python 3.6* programming language was used for the image acquisition via *picamera* package [35], which enables the capture of raw 10-bit Bayer images [36]. Adaptive integration times for individual LEDs (longer for the off-axis LEDs towards the edges of the array) enabled enhancement of the dynamic range and image signal-to-noise ratio. We chose to transfer all 256 images obtained by the microscope from the *Raspberry Pi 3* computer onto a desktop *Windows* computer to speed up the reconstruction. Reconstruction could also be performed on the *Raspberry Pi* with necessary optimization of recovery algorithms.

### Image reconstruction

Recorded images were demosaiced using bilinear interpolation from the *OpenCV* processing package [37] within the *Python 3.6* programming language. Before the reconstruction, the images were pre-processed by subtracting dark-frames to remove fixed pattern noise and all images were normalized according to their exposure times. The pre-processed images were divided into 128×128 pixel sub-images with an overlap of 28 pixels between adjacent image segments to aid in seamless stitching of the high-resolution reconstructions. Finally, LED-position calibration is performed independently on each image segment as described in the next section.

The FPM reconstruction algorithm is performed on each section of the FOV referred to as *I*^(*i*)^(**r**), where ***r*** is the coordinate vector in object space and *i* is the index corresponding to the LED used to illuminate and obtain the image. Before the reconstruction a high-resolution, wide-field object *o*(***r***) and its Fourier spectrum *O*(***k***) = ℱ[*o*(***r***)} are initialized by interpolating one of the low-resolution images to the required dimensions, where ***k*** is the coordinate vector in k-space and *ℱ* is the Fourier transform operator. The reconstruction steps described below are repeated for multiple iterations and within an *n*^th^ iteration, images obtained from illumination angles *i* are stitched together using the following steps:

1. Create a low-resolution target image Fourier spectrum estimate Ψ(***k***) by low-pass filtering the high-resolution, wide-field spectrum estimate with the pupil function *P*(***k***)

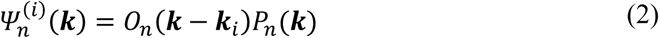

where ***k_i_*** is the k-space vector corresponding to angular LED illumination with an index *i*.
2. Create a low-resolution target estimate 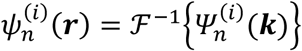 use it to create the updated low-resolution estimate 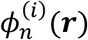 by replacing its amplitude with the experimentally obtained one

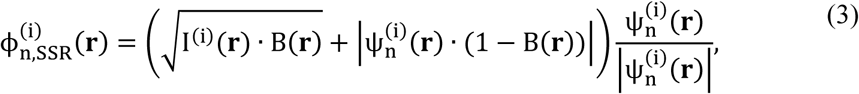

where B(**r**) is the binary Bayer matrix for the color channel being reconstructed. This is required if SSR is used [23], otherwise, if DR is being used then setting B(**r**) = 1 results in the standard amplitude update step

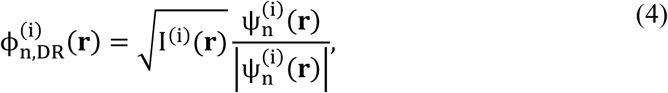
3. Create an updated low-resolution Fourier spectrum

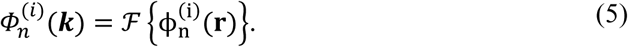
4. Update the high-resolution object Fourier spectrum *O(**k**)* using a second-order quasi Newton algorithm [38] together with embedded pupil recovery (EPRY) [16] and adaptive steps-size [39] schemes to improve convergence

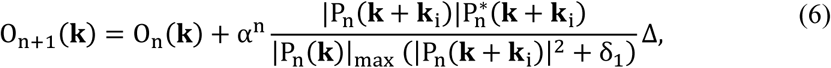

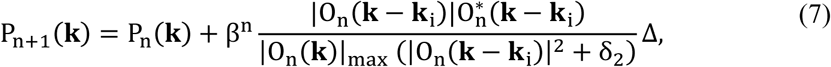

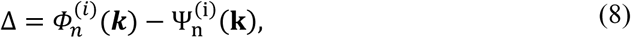

where *δ*_1_, *δ*_2_ are regularization parameters and *α*,*β* are adaptive-step size constants which are selected to improve convergence. More details on the pupil-aberration recovery framework are given in the following sections.

All reconstructed sections were stitched together to produce a full-FOV reconstructed image. Alignment and contrast variations were corrected prior to stitching. Histogram equalization with the central section is performed to remove contrast variations across the FOV for both amplitude and phase. Finally, all sections are blended together using *ImageJ* (using the *Fiji* plugin package) [40] to produce full-FOV images with seamless stitching.

All steps described above were performed for each of the red, green and blue channels independently and the final color image was assembled using linear image alignment with the scale-invariant feature transform (SIFT, part of the *Fiji* plugin package within *ImageJ)* [40] for each channel and mapping them into RGB color panes.

### Computational calibration of LED positions

An LED self-calibration method based on frequency-spectrum analysis of bright-field images [20] was used to locate pupil positions in spatial-frequency space for every 128×128 pixel section of the image, in order to accurately estimate the angle of illumination at the sample associated with each LED. A microscope objective acts as a low-pass filter and off-axis illumination shifts the frequencies in the object plane corresponding to the frequencies transmitted by the objective, enabling recording of higher spatial frequencies. These higher frequencies within the brightfield region appear as two overlapping circles in the Fourier transform of the intensity image, centered at the spatial frequency of the illumination angle. Finding the center of these circles yields the LED positions with sub-pixel accuracy, for every brightfield illumination angle [20] (Fig. 5(a)). After finding position displacements for each bright-field LED, a homographic transformation matrix that best represents the misalignment of the LED array is derived. This transformation matrix is applied to dark-field LEDs as well. However, non-linear distortions, such as field curvature [25], make LED positions appear to be distorted differently across the FOV. To mitigate this problem, we split the full FOV image into 128×128 pixel sections and apply LED calibration for each section individually. If non-linear distortions are present, then each section will have a different LED array translation shown in Fig. 5(c). These distortions were corrected using an affine transformation that best represents corrections for each section of the FOV.

**Figure 5.**
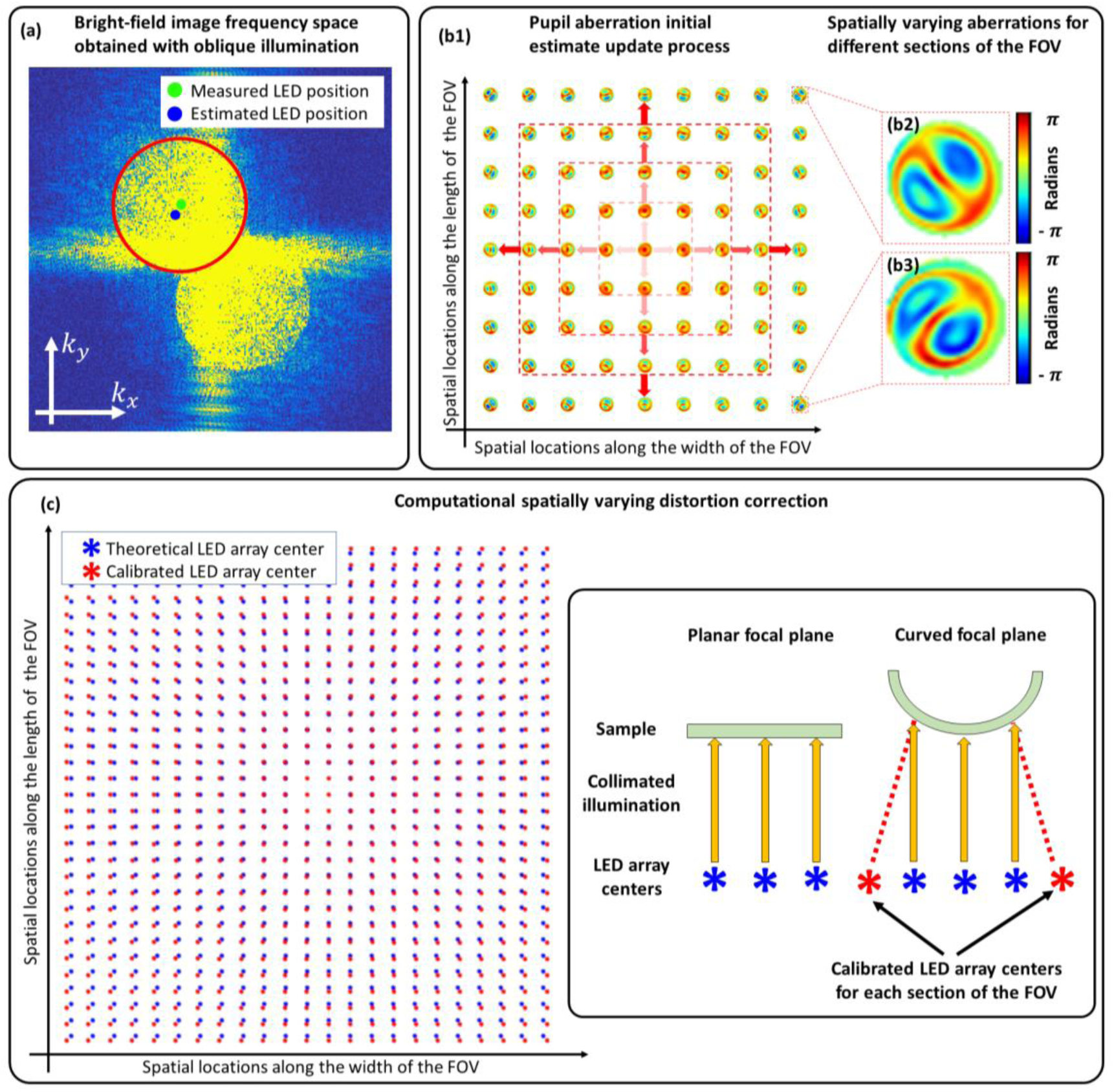
(**a**) Frequency space of a brightfield image obtained using an oblique illumination angle. Blue and green dots indicate initial and corrected LED positions respectively. (**b1**) Aberrations recovered from each section are used as initial estimates for neighboring sections, starting from the center of the FOV towards the edges. (**b2, b3**) Examples of recovered aberrations throughout the full-FOV indicating spatially-varying aberrations. (**c**) Implementing LED calibration on each segment across the FOV enabled us to find the spatially varying distortion by measuring the global LED position shift.

### Computational aberration correction

Spatially-varying aberrations for each segment of the FOV are recovered using the EPRY algorithm [16] to enable FPM reconstruction of the images. However, our microscope suffers from aberrations that increase progressively towards the edges of the FOV, and the EPRY algorithm fails for the more highly aberrated sections. A good initial estimate of the aberrations is required for the EPRY algorithm to converge. Therefore, starting with the central 128×128 section of the FOV, we run the EPRY recovery step for 40 iterations, reset the recovered image intensity and phase while retaining the aberrations, and iterate the algorithm for 3 more times. The reset step forces the algorithm to escape from local minima and enables convergence towards a global solution. We use the recovered central aberrations as an initial estimate for the surrounding sections (Fig. 5(b)). This update process continues until aberrations for every section of the FOV are recovered.

Low-cost lenses, such as the ones we have used, tend to suffer from severe chromatic aberrations. We found that when the microscope is focused using one color of LED, the chromatic aberration (primarily defocus) for images recorded using other colors was significant to cause the reconstruction algorithms to fail. The aberrations recovered from the central section of the color where the microscope is focused are used as an initial estimate for the defocused color that is being processed. This involved decomposition of recovered pupil aberrations into 30 Zernike coefficients using the singular value decomposition function in *MATLAB* from which the chromatically-aberrated pupil functions were estimated.

## Supplementary material 1: Instructions to build a *Raspberry Pi* Fourier ptychographic computational microscope

This document provides instructions to build a low-cost computational microscope reported in the manuscript: “*Low-cost, sub-micron resolution, wide-field computational microscopy with Raspberry Pi hardware*”. The CAD files and data acquisition codes can be downloaded from http://dx.doi.org/10.5525/gla.researchdata.594.

### Introduction

One of the aims when building this microscope was to use only off-the-shelf components that can be easily bought anywhere and to design the microscope in such a way that it could be assembled with minimal external components. Avoiding complexity allowed us to build a very low-cost and robust microscope, which can be assembled and used with opensource software. Designs for the parts were made using *OpenSCAD* open source CAD software and printed with *Ultimaker 2+* 3D printer.

The microscope was designed around the *<u>Raspberry Pi 3</u>* computer board due to a wide opensource community and the support available. The computer itself has a CSI port to which a *<u>Raspberry Pi</u>* camera can be connected. For the illumination we used a *<u>Unicorn HAT HD</u>* 16×16 LED array, which is an add-on designed for the *Raspberry Pi* boards. It mounts directly onto the GPIO pins on top of the board. Camera and the LED board can be connected and controlled easily via opensource libraries available for *Python* or C++ programming languages.

Furthermore, *Raspberry Pi* camera comes mounted with a mobile-phone-camera type lens. It was unscrewed from the camera and used as our microscope objective. The component list required to build the setup is provided below, along with a step-by-step instruction set for assembly and operation of the microscope.

### Component list

**Supplementary Table S1 1.** List of the off-the-shelf components required to build the microscope.

| Off-the-shelf components | Quantity | Purpose |
| --- | --- | --- |
| Raspberry Pi 3 computer board | 1 | Controlling the camera and LED array;<br>image capture and storage |
| 20mm long M3 screws | 4 | Fix Raspberry Pi computer board to a<br>stable base |
| Unicorn HAT HD 16x16 LED array | 1 | Illumination source |
| Raspberry Pi V2.1 NoIR camera | 1 | Image sensor |
| Unscrewed lens from the Raspberry<br>Pi camera (Part 4) | 1 | Microscope objective |
| 3.5mm diameter, >10mm long,<br>0.25mm pitch screw and bushing<br>(Thorlabs F3ES25, F3ESN1P) | 1 | Used for high-accuracy focusing |
| >40mm long M6 screws + nuts | 2 | Attaching the focusing stage to the<br>sample stage |
| 5.6mm diameter, ~10mm long springs<br>(RS Stock No. 821-431) | 2 | Counter balance force for the focusing<br>stage |
| 5mm long M1.5 screws | 4 | Screwing *Raspberry Pi* (part 4) camera<br>to the focusing stage |

### Design

**Supplementary Figure S1 1.**
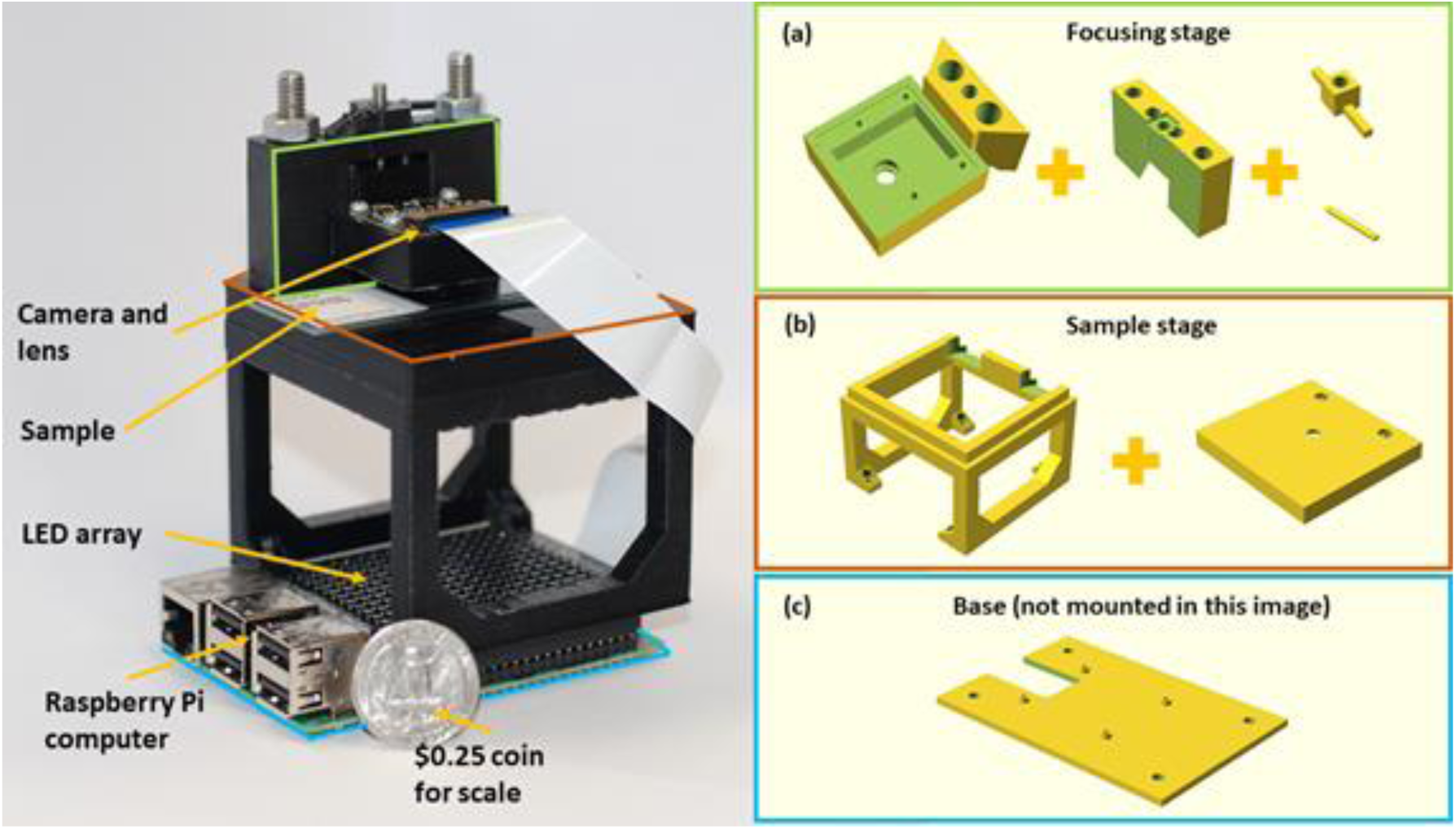
The experimental setup with all the necessary annotations for reference to 3D printed designs.

Supplementary Figure S1 1 shows the assembled setup together with 3D printed parts required. It was built using components described in Supplementary Table S1 1. Once each component is 3D printed, the assembly is very simple and requires only a few screwdrivers.

#### Base

The plastic base shown in Supplementary Figure S1 1(c) was printed such that the Raspberry pi and the sample stage could be screwed onto it. The base itself has 4 holes which can be used for screwing the microscopes to the optical bench if needed. This was designed to provide higher stability when longitudinal imaging might be required.

#### Sample stage

The sample stage shown in Supplementary Figure S1 1(b) was designed to be mounted on the Raspberry Pi board with an LED array on top. The four screw holes on the 3D printed sample stage match those found on the Raspberry Pi and the plastic base. All components can be screwed tightly together to form a single microscope unit.

There is another 3D printed part that goes on top of the sample stage legs. It has 3 holes on it where the central one acts as an aperture for the sample, reducing any stray light and reflections from the LED array. The other 2 holes were made for screws that attach the focusing stage to the sample stage.

#### Camera holder and focusing stage

The focusing stage shown in Supplementary Figure S1 1(a) is composed of four 3D printed elements.

The camera holder module (the first 3D printed part seen in Supplementary Figure S1 1(a)) was designed to mount the microscope objective (the unscrewed camera lens) in place and screw the camera above it. This was designed for finite-conjugate microscope configuration to be established. The unscrewed lens from the camera has a 1.5mm aperture only on one side; lens must be mounted such that the aperture is facing downwards (towards the sample). This compact design was set to achieve 1.5× magnification, but it can be easily modified by changing the distance between the lens and the detector.

The camera holder mount, (the second 3D printed part seen in Supplementary Figure S1 1(a)) serves several purposes including focusing the sample. Firstly, it has rails onto which camera holder module is mounted and can be moved up or down for focusing. The central hole in the camera holder mount is for the 0.25mm pitch screw. Springs are fed through the inner pair of holes in the camera holder mount and the corresponding holes in the camera holder module. They are held in place by sliding the pins shown in Supplementary Figure S1 1(a) through each end of both springs. The screw is used to push down on the camera holder module while the springs and bottom pin provide a counter force to push it upwards. This way the module can slide along the rails with high-precision, by turning the screw. Springs provide stability and push the module upwards when the screw does not provide a downward force anymore, which should minimize the backslash error.

Secondly, the outer holes in the camera holder mount enables addition of screws or bolts to attach the whole focusing module to the top of the sample stage. While the focusing is done via a translation stage, the sample must be translated by hand. In our setup, the FOV is large so precise translation is not required; hence, we chose to use this design. However, there are 3D printed sample translation stages available in the opensource community that can be integrated into our design.

### Assembly instructions

Access to a 3D printer is required to print several parts required for the assembly. We used *Ultimaker 2+* with a nozzle size of 0.25mm for the camera holder module and 0.4mm for the other components. Also, the lens from the *Raspberry Pi V2.0* camera must be unscrewed before the assembly. Step-by-step instructions to assemble the microscope:

1. 3D print all the parts using a printer of your choice. We used *openSCAD* to design, render and save the designs in .STL format. *CURA* software was used to create the files that can be read by the *Ultimaker 2+* 3D printer. Black PLA filament and a 0.4mm diameter nozzle was used for printing the sample stage parts, while a 0.25mm nozzle was used to print the focusing stage. Our files were designed to match the tolerance of the nozzles on our printer. The 3D models need to be tweaked when a different nozzle size or a different 3D printer is used due to change in the tolerances.
2. Connect the *Raspberry Pi* camera to the *Raspberry Pi* board.

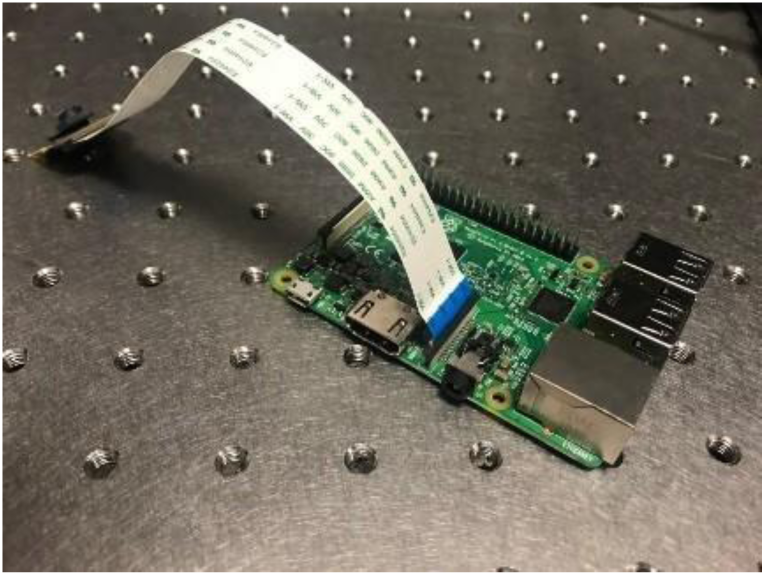
3. Mount the LED array on top of the Raspberry Pi board by plugging it into the GPIO pins on the board.

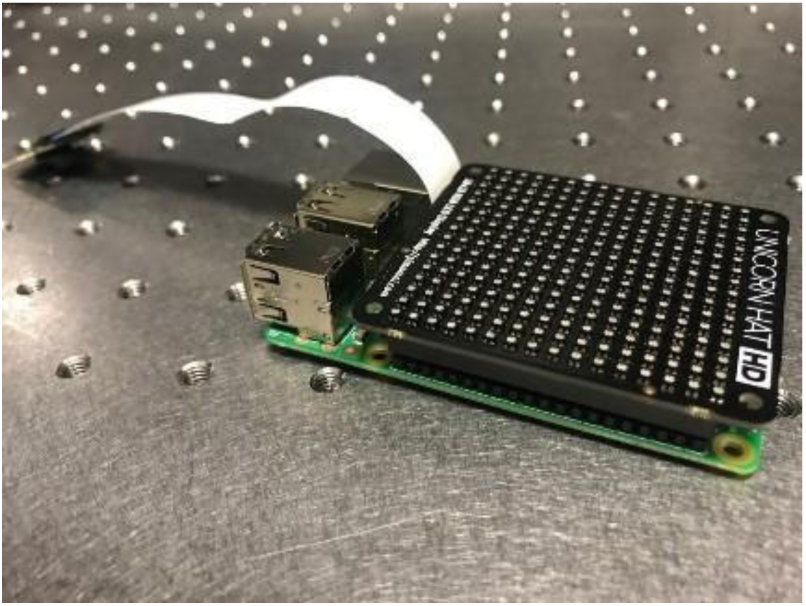
4. Place the sample stage such that the screw holes match the base; making sure that the sample-stage feet are not on top of any of the LEDs. Then place a nut in each foot of the sample stage.
5. Place the spacers in between the *Raspberry Pi* board *UnicornhatHD* board so that they are aligned with the screw holes and then screw the *Raspberry Pi* board and the sample stage to the base such that they form a single rigid module.

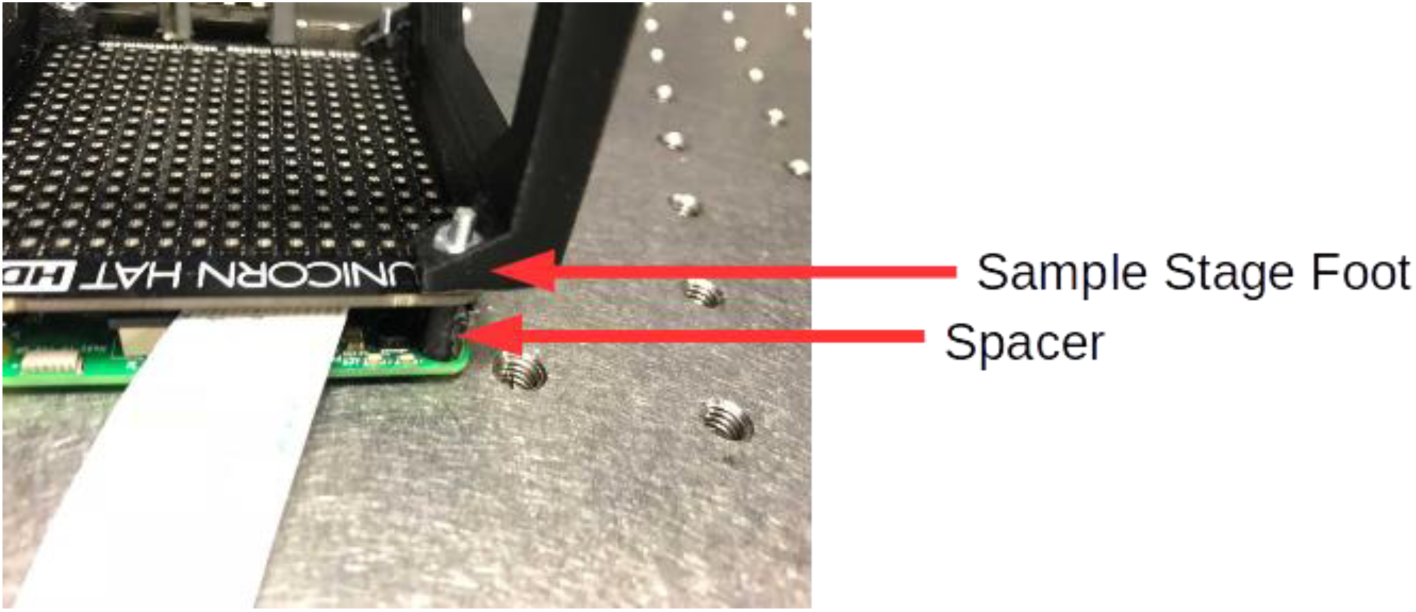
6. Take the camera holder; place the lens in the circular slot with the aperture facing downwards, towards the sample stage.

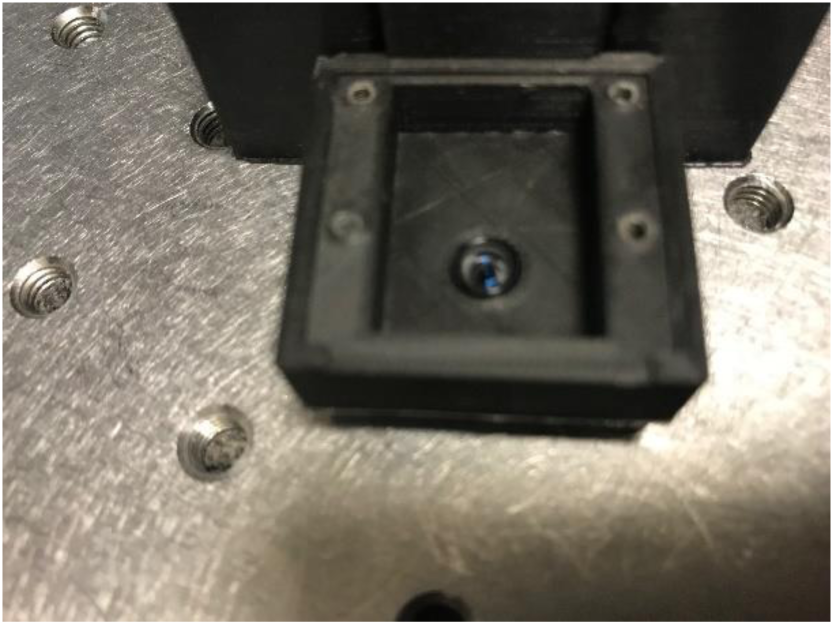
7. Mount the camera, align with the screw holes of the 3D printed camera holder; screw it in place tightly.

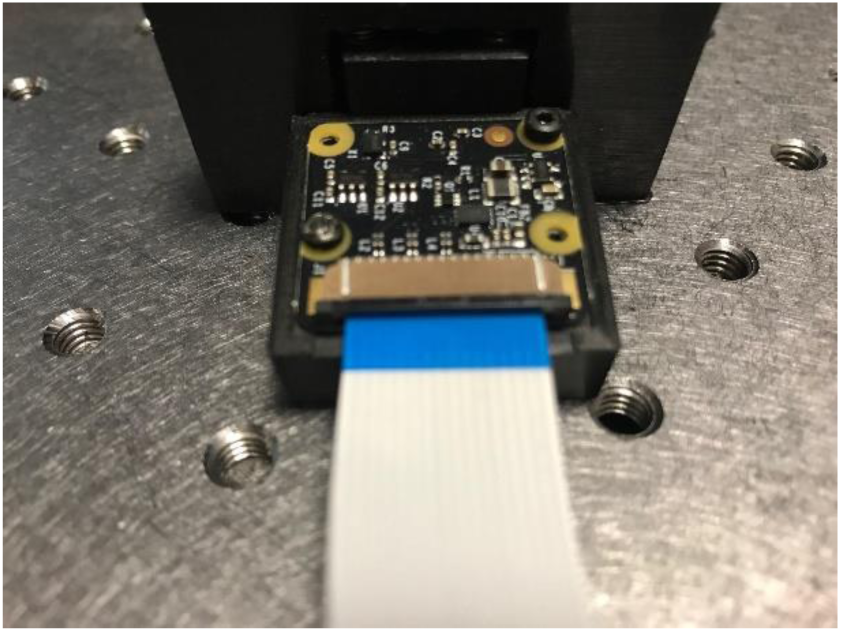
8. Slide the camera holder module onto the focusing stage rails. Thread the springs through the holes on the focusing stage and the camera module as shown by a red arrow in the figure below. Use two 3D printed horizontal pins seen in Supplementary Figure S1 1(a)) to hold the ends of the springs at the top and bottom of the focusing stage; the spring should be long enough such that it is stretched out and apply a strong counter-balance force to the screw.

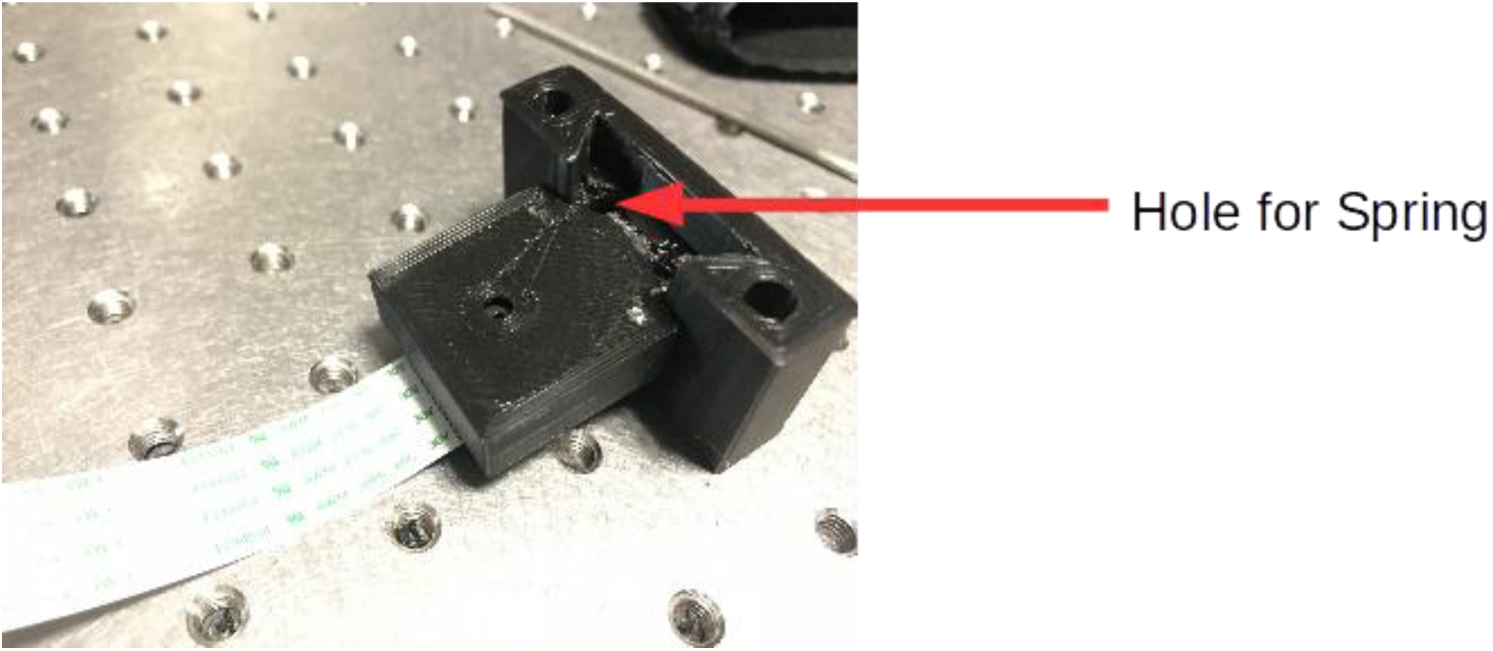

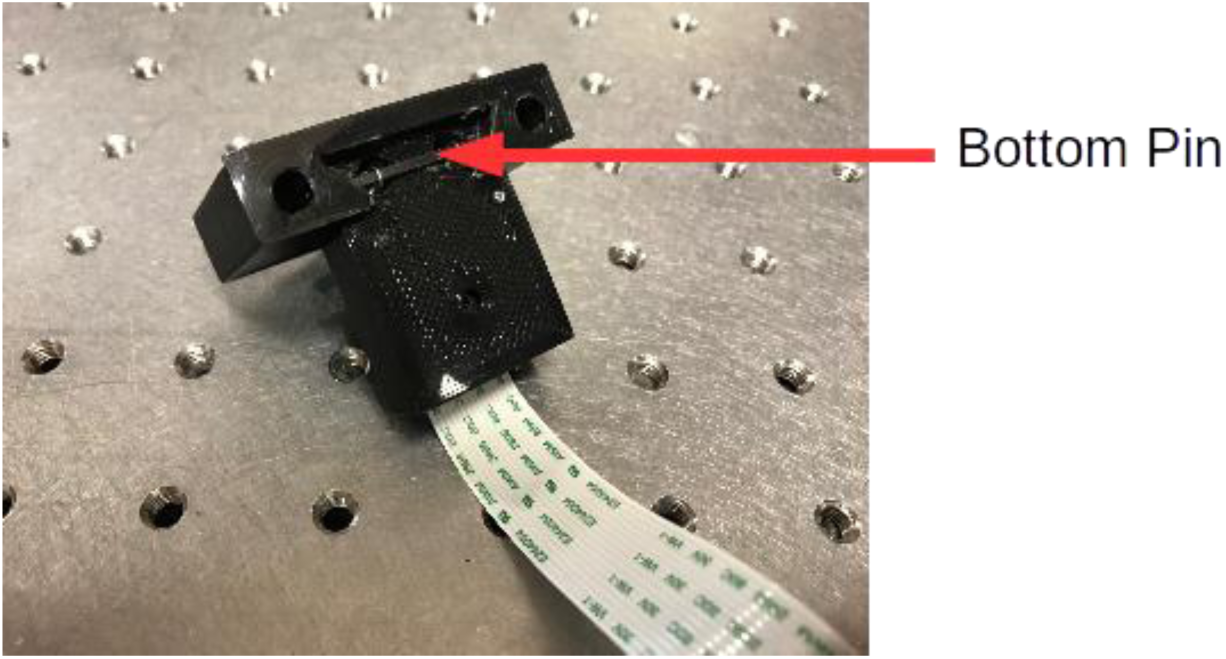
9. Pull down the camera holder module and, from underneath, place the bushing up into the central hole of the focusing stage. Then, place the screw in the top of the camera holder mount. Screw it in such that the screw pushes onto the camera holder module.
10. Use screws with nuts to fix the focusing stage tightly to the top plate of the sample stage from Supplementary Figure S1 1(b)).
11. Place the focusing stage module onto the sample stage module with the *Raspberry Pi* computer. The part is designed to have a tight fit, if it is not tight, please adjust the tolerances.
12. Optional: connect a screen using the HDMI port.
13. Optional: connect a keyboard and a mouse.
14. Optional: Place the *Raspberry Pi* board on top of the 3D printed base; align the base and the Raspberry Pi such that the screw holes are on top of each other.

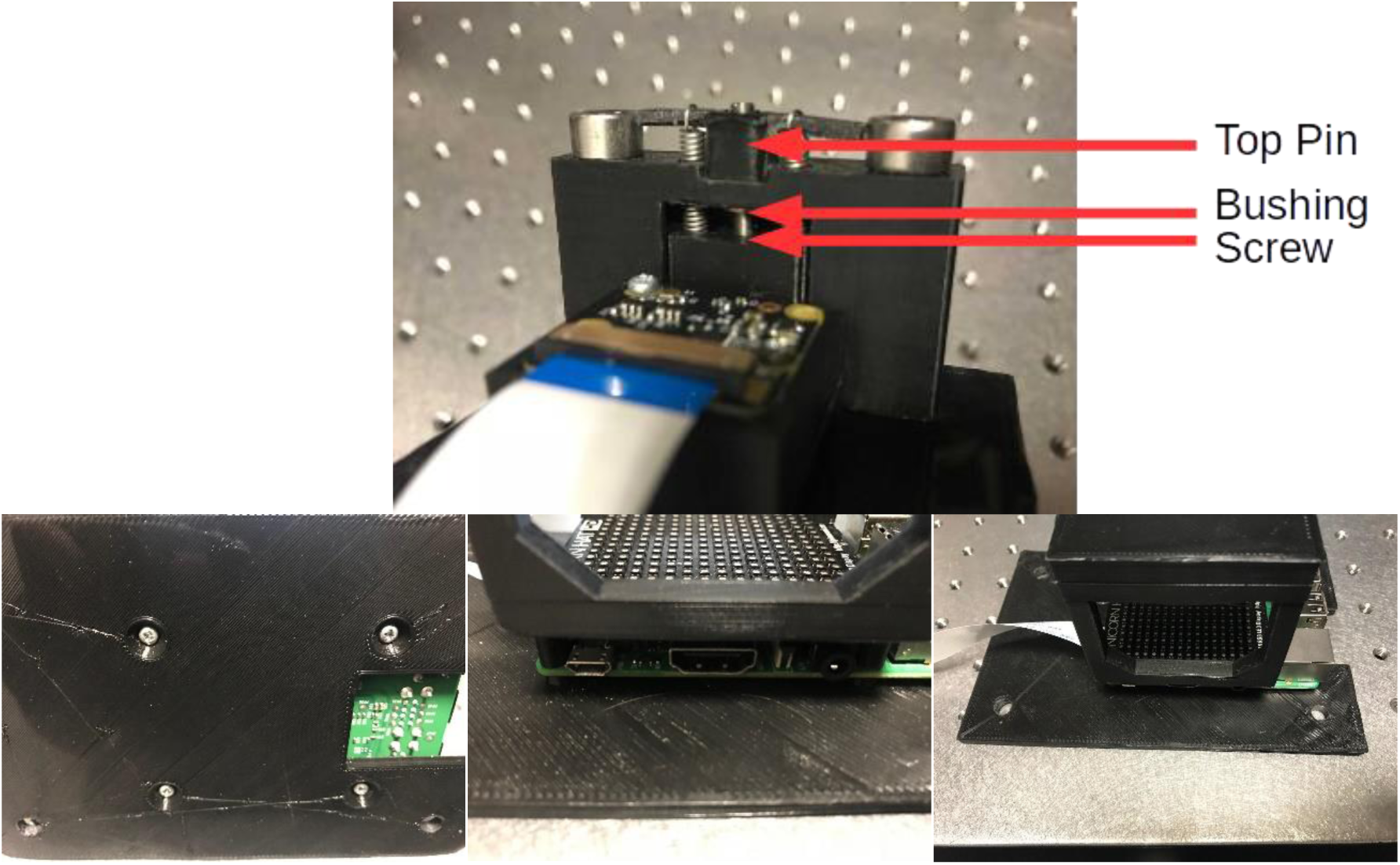

### Operating the Microscope

#### Installing Software on the Raspberry Pi

Raspbian is a free *Raspberry Pi* operating system available for download from the manufacturer’s website. It can be installed by following the guide listed in the *Links* section.

The various interfaces of the Raspberry Pi can be enabled by going to Applications Menu -> Preferences -> Raspberry Pi Configuration -> Interfaces, and then enabling all options.

Image acquisition codes can also be downloaded from the *Links* section. Various python packages will need to be installed before these can be used. The packages needed are:

- -Unicornhathd
- -Numpy
- -Picamera
- -Matplotlib
- -Io
- -Random
- -Fractions

These can be installed using the pip package management system or by installing anaconda on the raspberry pi. However, the <u>picamera</u> and <u>unicornhathd</u> packages are not included in anaconda and will need to be installed separately. Links to the installation guides of these packages and a more <u>general guide</u> to installing python packages on the Raspberry Pi are provided in the *Links* section. Python2 is used for image acquisition so follow instructions for Python2.7 as opposed to Python3.

#### Data Acquisition

1. Connect the Raspberry Pi to a keyboard, mouse and monitor, and turn it on.
2. Place the sample on the top plate of the sample stage, underneath the camera mount.
3. Use the “Focusing” script to make sure the sample is positioned correctly and in focus. This script has an option to zoom that can be used if needed. To focus the microscope, use an Allen key to turn the screw in the focusing stage.
4. Close the preview and open the “main data acquisition” file.
5. Adjust the necessary parameters in the data acquisition file and save the file.
6. Place the microscope in a dark room or cover it, being careful to ensure the sample is not moved and the focus is not shifted.
7. Run the data acquisition script
8. The captured data can be copied by a USB drive or the SD card on the *Pi* can be inserted into a PC and *<u>disk internals Linux reader</u>* can be used to copy the data.
9. Switch off the Raspberry Pi after data acquisition is complete.

### Links

- Data acquisition codes and CAD files: http://dx.doi.org/10.5525/gla.researchdata.594
- *Raspbian* installation guide: https://www.raspberrypi.org/documentation/installation/installing-images/README.md
  - Needs Etcher software to install Raspbian on the SD Card
  - But first, download the Raspbian image from the above link.
- Guide to Installing *Python Packages* on *Raspberry Pi: https://www.raspberrvpi.org/documentation/linux/software/python.md*
- *Picamera* installation guide: http://www.picamera.readthedocs.io/en/release-1.0/install2.html
- *UnicornHatHD* installation guide: https://www.github.com/pimoroni/unicorn-hat-hd
- *Pimoroni Unicorn HD* LED array: https://shop.pimoroni.com/products/unicorn-hat-hd
- *Raspberry Pi V2.1* camera: https://www.raspberrypi.org/products/camera-module-v2/
- *Disklnternals Linux Reader* can be used to read the files on a Linux SD card from a computer: https://www.diskinternals.com/linux-reader/

## Supplementary material 2: Reconstructions from the simulations

Simulations were carried out to compare the reconstruction quality from the sparsely sampled Bayer filtered images using (1) standard FPM algorithms on demosaiced images and (2) sparsely sampled FPM reconstruction on raw Bayer data. These reconstruction methods were applied to investigate their performance for various sampling and frequency overlap criteria. Simulations for a non-Bayer image sensor (monochrome) are also presented to provide a reference for the results obtained from a Bayer filtered image sensor (colour). Results are shown in Supplementary Figure S2 1, Supplementary Figure S2 2, Supplementary Figure S2 3. These are the reconstructed images for data points displayed in the graphs presented in Figure 2 of the main manuscript.

**Supplementary Figure S2 1.**
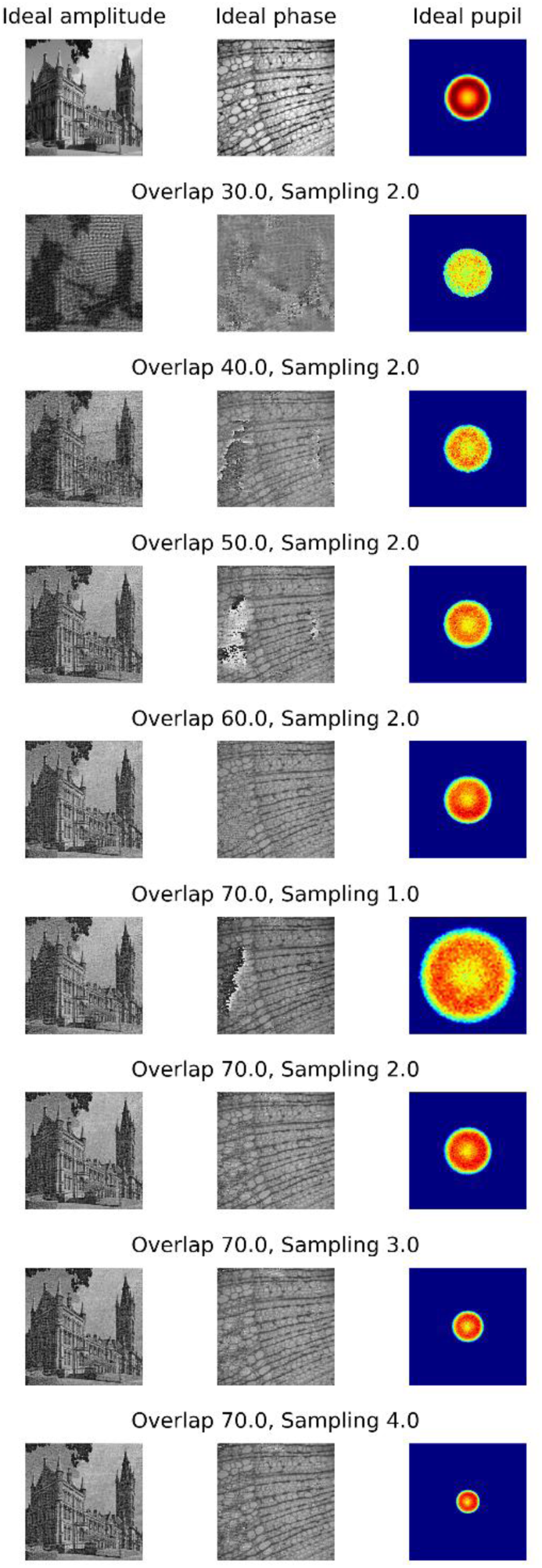
Reconstructions from images obtained with a monochrome sensor (no Bayer filter) using the standard FPM algorithm. First row shows the expected ideal reconstruction and the remaining rows shows the reconstructions from datasets captured with (1) various image sampling criteria and (2) overlap between the spatial frequencies captured by any two adjacent illumination angles. Noise and aberrations are added in the simulated images to mimic the experimental conditions.

**Supplementary Figure S2 2.**
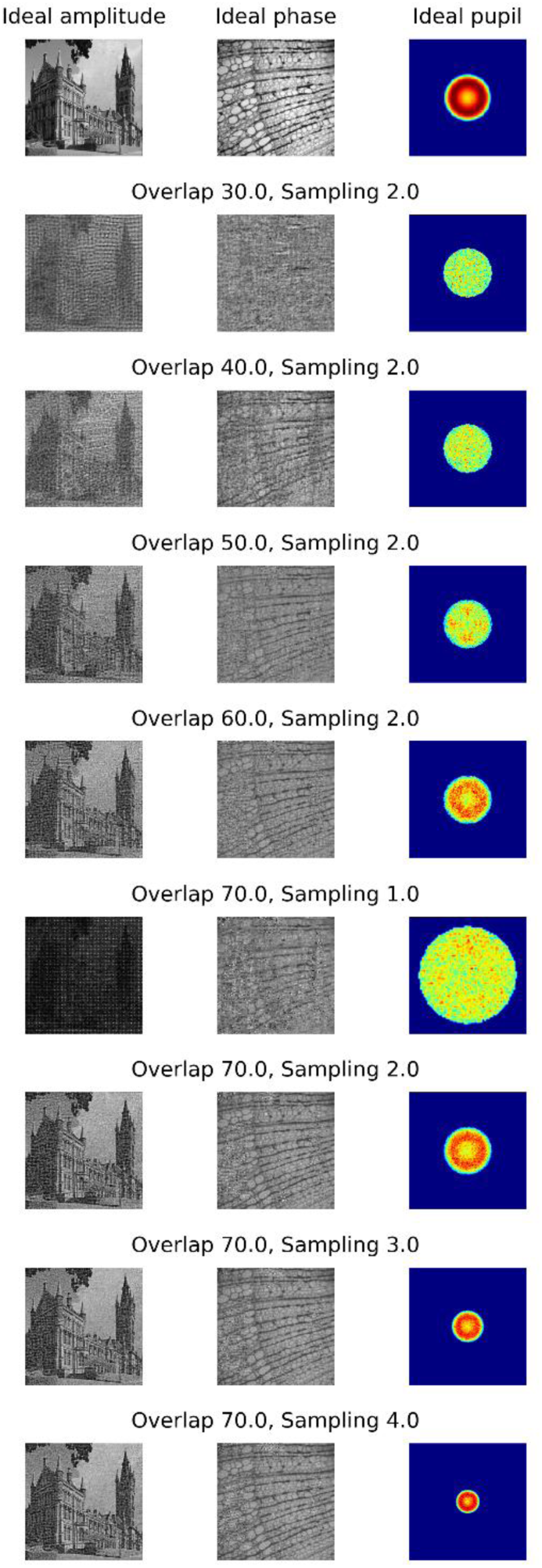
Reconstructions from images obtained with a color sensor (with Bayer filter array) using the sparsely-sampled FPM algorithm. These images are sparse due to the intermittent sampling from the Bayer filter array. First row shows the expected ideal reconstruction and the remaining rows shows the reconstructions from datasets captured with (1) various image sampling criteria and (2) overlap between the spatial frequencies captured by any two adjacent illumination angles. Noise and aberrations are added in the simulated images to mimic the experimental conditions.

**Supplementary Figure S2 3.**
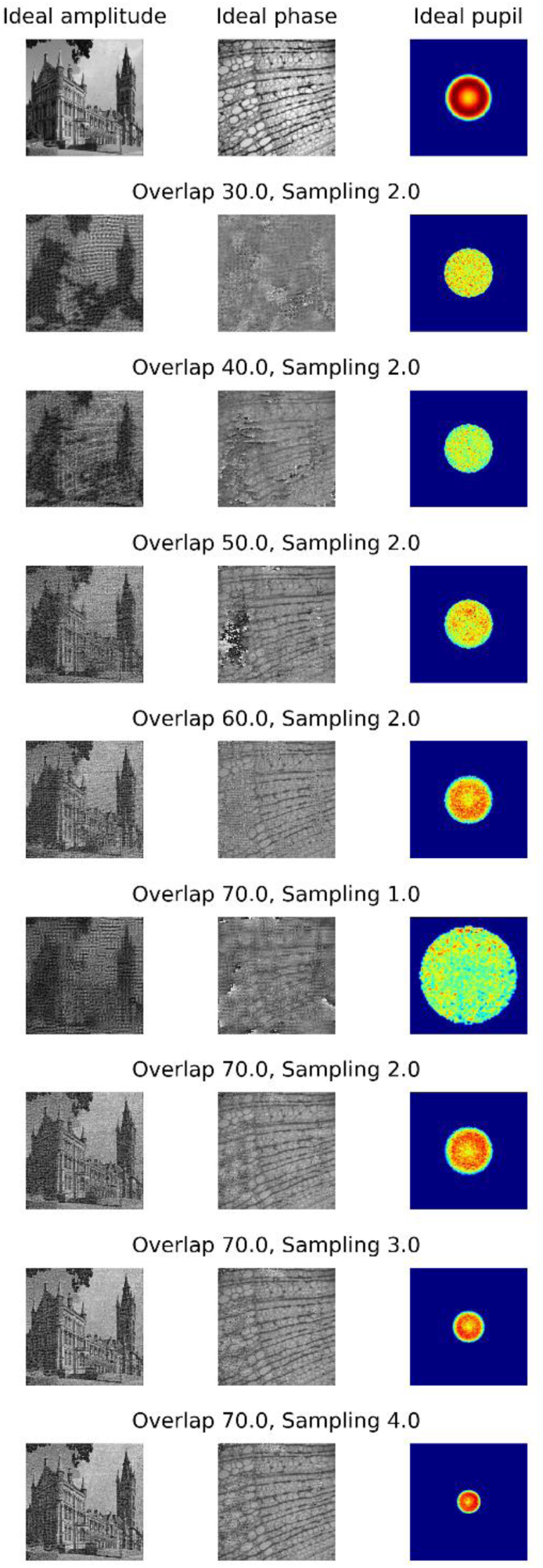
Reconstructions from images obtained with a color sensor (with Bayer filter array) using the standard FPM algorithm after demosaicing. The sparse images captured from the Bayer filter array are demosaiced (bilinear interpolation) such that the standard FPM algorithm can be implemented. First row shows the expected ideal reconstruction and the remaining rows shows the reconstructions from datasets captured with (1) various image sampling criteria and (2) overlap between the spatial frequencies captured by any two adjacent illumination angles. Noise and aberrations are added in the simulated images to mimic the experimental conditions.

## Acknowledgments

We thank Victor Lovic for his help in 3D printing and building the setup, Nicholas Nugent for his help in writing the tutorial and we are grateful to Dr. Jonathan Taylor for his feedback on the manuscript.

## Author contributions

P.C.K conceived the idea, contributed to the design, development of the algorithms and theory. P.C.K and A.H oversaw the experiments. T.A designed and performed the experiments, developed the final reconstruction and calibration algorithms and contributed to the development of the theory. R.E and L.W provided code and support for misalignment calibration. All authors participated in the writing and the revisions of the manuscript.

## Data and materials availability

Raw data and computer codes used for image processing can be obtained from http://dx.doi.org/10.5525/gla.researchdata.687. Data acquisition codes and 3D-printed designs can be obtained from http://dx.doi.org/10.5525/gla.researchdata.594.

## References

1. C. A. Petti, C. R. Polage, T. C. Quinn, A. R. Ronald, and M. A. Sande, “Laboratory Medicine in Africa: A Barrier to Effective Health Care,” Clin. Infect. Dis. 42, 377–382 (2006).

2. M. Imwong, S. Hanchana, B. Malleret, L. Rénia, N. P. J. Day, A. Dondorp, F. Nosten, G. Snounou, and N. J. White, “High-throughput ultrasensitive molecular techniques for quantifying low-density malaria parasitemias,” J. Clin. Microbiol. 52, 3303–9 (2014).

3. D. Mendlovic, A. W. Lohmann, and Z. Zalevsky, “Space–bandwidth product adaptation and its application to superresolution: examples,” J. Opt. Soc. Am. A 14, 563 (1997).

4. P. C. Konda, “Multi-Aperture Fourier Ptychographic Microscopy : Development of a high-speed gigapixel coherent computational microscope,” (2017).

5. G. McConnell, J. Trägårdh, R. Amor, J. Dempster, E. Reid, and W. B. Amos, “A novel optical microscope for imaging large embryos and tissue volumes with sub-cellular resolution throughout,” Elife 5, 1–15 (2016).

6. G. Zheng, X. Ou, and C. Yang, “0.5 Gigapixel Microscopy Using a Flatbed Scanner,” Biomed. Opt. Express 5, 1–8 (2013).

7. D. N. Breslauer, R. N. Maamari, N. A. Switz, W. A. Lam, and D. A. Fletcher, “Mobile phone based clinical microscopy for global health applications,” PLoS One 4, 1–7 (2009).

8. N. A. Switz, M. V. D’Ambrosio, and D. A. Fletcher, “Low-cost mobile phone microscopy with a reversed mobile phone camera lens,” PLoS One 9, (2014).

9. Z. J. Smith, K. Chu, A. R. Espenson, M. Rahimzadeh, A. Gryshuk, M. Molinaro, D. M. Dwyre, S. Lane, D. Matthews, and S. Wachsmann-Hogiu, “Cell-phone-based platform for biomedical device development and education applications,” PLoS One 6, (2011).

10. Z. F. Phillips, M. V. D’Ambrosio, L. Tian, J. J. Rulison, H. S. Patel, N. Sadras, A. V. Gande, N. A. Switz, D. A. Fletcher, and L. Waller, “Multi-contrast imaging and digital refocusing on a mobile microscope with a domed LED array,” PLoS One 10, 1–13 (2015).

11. J. P. Sharkey, D. C. W. Foo, A. Kabla, J. J. Baumberg, and R. W. Bowman, “A one-piece 3D printed flexure translation stage for open-source microscopy,” Rev. Sci. Instrum. 87, (2016).

12. A. Skandarajah, C. D. Reber, N. A. Switz, and D. A. Fletcher, “Quantitative imaging with a mobile phone microscope,” PLoS One 9, (2014).

13. J. S. Cybulski, J. Clements, and M. Prakash, “Foldscope: Origami-based paper microscope,” PLoS One 9, (2014).

14. G. Zheng, R. Horstmeyer, and C. Yang, “Wide-field, high-resolution Fourier ptychographic microscopy,” Nat. Photonics 7, 739–745 (2013).

15. J. W. Goodman, Introduction to Fourier Optics (2005), Vol. 8.

16. X. Ou, G. Zheng, and C. Yang, “Embedded pupil function recovery for Fourier ptychographic microscopy,” Opt. Express 22, 4960–72 (2014).

17. L.-H. Yeh, L. Tian, Z. Liu, M. Chen, J. Zhong, and L. Waller, “Experimental robustness of Fourier Ptychographic phase retrieval algorithms,” Imaging Appl. Opt. 2015 23, CW4E.2 (2015).

18. L. Tian, Z. Liu, L.-H. Yeh, M. Chen, J. Zhong, and L. Waller, “Computational illumination for high-speed in vitro Fourier ptychographic microscopy,” Optica 2, 904–911 (2015).

19. Z. Bian, S. Dong, and G. Zheng, “Adaptive system correction for robust Fourier ptychographic imaging,” Opt. Express 21, 32400–10 (2013).

20. R. Eckert, Z. F. Phillips, and L. Waller, “Efficient illumination angle self-calibration in Fourier ptychography,” Appl. Opt. 57, 5434 (2018).

21. S. Dong, K. Guo, P. Nanda, R. Shiradkar, and G. Zheng, “FPscope: a field-portable high-resolution microscope using a cellphone lens,” Biomed. Opt. Express 5, 3305–10 (2014).

22. M. Pagnutti, R. E. Ryan, G. Cazenavette, M. Gold, R. Harlan, E. Leggett, and J. Pagnutti, “Laying the foundation to use Raspberry Pi 3 V2 camera module imagery for scientific and engineering purposes,” J. Electron. Imaging 26, 013014 (2017).

23. S. Dong, Z. Bian, R. Shiradkar, and G. Zheng, “Sparsely sampled Fourier ptychography,” Opt. Express 22, 5455 (2014).

24. J. Sun, Q. Chen, Y. Zhang, and C. Zuo, “Sampling criteria for Fourier ptychographic microscopy in object space and frequency space,” Opt. Express 24, 15765 (2016).

25. L. Waller and L. Tian, “3D Phase Retrieval with Computational Illumination,” in Imaging and Applied Optics 2015, OSA Technical Digest (Online) (Optical Society of America, 2015), p. CW4E.1.

26. K. Guo, S. Dong, and G. Zheng, “Fourier Ptychography for Brightfield, Phase, Darkfield, Reflective, Multi-Slice, and Fluorescence Imaging,” IEEE J. Sel. Top. Quantam Electron. 22, 1–12 (2016).

27. Z. Liu, L. Tian, S. Liu, and L. Waller, “Real-time brightfield, darkfield, and phase contrast imaging in a light-emitting diode array microscope,” J. Biomed. Opt. 19, 106002 (2014).

28. S. Dong, R. Shiradkar, P. Nanda, and G. Zheng, “Spectral multiplexing and coherent-state decomposition in Fourier ptychographic imaging,” Biomed. Opt. Express 5, 22817–22825 (2014).

29. L. Tian, X. Li, K. Ramchandran, and L. Waller, “Multiplexed coded illumination for Fourier Ptychography with an LED array microscope,” Biomed. Opt. Express 162, 4960–4972 (2014).

30. T. Nguyen, Y. Xue, Y. Li, L. Tian, and G. Nehmetallah, “Convolutional neural network for Fourier ptychography video reconstruction: learning temporal dynamics from spatial ensembles,” (2018).

31. J. V. Tu, “Advantages and disadvantages of using artificial neural networks versus logistic regression for predicting medical outcomes,” J. Clin. Epidemiol. 49, 1225–1231 (1996).

32. P. C. Konda, J. M. Taylor, and A. R. Harvey, “Scheimpflug multi-aperture Fourier ptychography: coherent computational microscope with gigapixels/s data acquisition rates using 3D printed components,” in High-Speed Biomedical Imaging and Spectroscopy: Toward Big Data Instrumentation and Management II (2017), Vol. 10076, p. 100760R.

33. P. C. Konda, J. M. Taylor, and A. R. Harvey, “Parallelized aperture synthesis using multi-aperture Fourier ptychographic microscopy,” arXiv Prepr. arXiv ID 1806.02317 (2018).

34. Z. F. Phillips, R. Eckert, and L. Waller, “Quasi-Dome : A Self-Calibrated High-NA LED Illuminator for Fourier Ptychography,” in Imaging and Applied Optics 2017 (2017).

35. D. Jones, “Picamera 1.13 Documentation,” https://picamera.readthedocs.io/en/release-1.13/.

36. S. van der Walt, S. C. Colbert, and G. Varoquaux, “The NumPy Array: A Structure for Efficient Numerical Computation,” Comput. Sci. Eng. 13, 22–30 (2011).

37. G. R. Bradski and A. Kaehler, Learning OpenCV: Computer Vision with the OpenCV Library (O’Reilly, 2008).

38. L.-H. Yeh, J. Dong, J. Zhong, L. Tian, M. Chen, G. Tang, M. Soltanolkotabi, and L. Waller, “Experimental robustness of Fourier Ptychography phase retrieval algorithms,” Opt. Express 23, 38–43 (2015).

39. C. Zuo, J. Sun, and Q. Chen, “Adaptive step-size strategy for noise-robust Fourier ptychographic microscopy,” Opt. Express 24, 4960–4972 (2016).

40. J. Schindelin, I. Arganda-Carreras, E. Frise, V. Kaynig, M. Longair, T. Pietzsch, S. Preibisch, C. Rueden, S. Saalfeld, B. Schmid, J.-Y. Tinevez, D. J. White, V. Hartenstein, K. Eliceiri, P. Tomancak, and A. Cardona, “Fiji: an open-source platform for biological-image analysis,” Nat. Methods 9, 676–682 (2012).

